# Diammonium glycyrrhizinate injection promotes zebrafish angiogenesis through the mTOR/HIF-1 signaling pathway

**DOI:** 10.1101/2025.02.19.639026

**Authors:** Shuqing Yu, Jiahao Yu, Fengzhi Sun, Xuanming Zhang, Wenlong Sheng, Kechun Liu, Xiaobin Li, Wei Li

**Affiliations:** Engineering Research Center of Zebrafish Models for Human Diseases and Drug Screening of Shandong Province Biology Institute, Qilu University of Technology (Shandong Academy of Sciences) 28789 East Jingshi Road Jinan 250103, Shandong, P.R. China; Department of Pharmacy, Nanjing Drum Tower Hospital Group Suqian Hospital, Department of Pharmacy,Suqian Hospital Affiliated to Xuzhou Medical University, Xuzhou 221004, Jiangsu, China

**Keywords:** Glycyrrhizic acid preparation, Diammonium glycyrrhizinate injection, Drug screening, Zebrafish, Pro-angiogenesis, mTOR/HIF-1 signaling pathway

## Abstract

**Background:** Glycyrrhizic acid (GA) is the main component of the traditional Chinese medicine Glycyrrhiza glabra. Many GA preparations in the pharmaceutical market have good efficacy in hepatoprotection and the treatment of liver diseases. However, the pro-angiogenic effects for GA-based agents have not been researched specifically. The aim of this study was to investigate the pro-angiogenic activity of GA preparations commonly used in clinical practice and screen the representative angiogenesis promotor to further elucidate the mechanism of action

**Methods:** A zebrafish vascular injury model induced by PTK787 was used to assess the pro-angiogenic effect of GA preparations. The pro-angiogenic activity of diammonium glycyrrhizinate injection (DGI) was further systematically evaluated by detecting the growth of the intersegmental vessels (ISVs) and subintestinal vein vessels (SIVs) in zebrafish. The underlying mechanism of action for DGI was explored by transcriptomics analysis and RT-qPCR validation;

**Results:** The results showed that magnesium isoglycyrrhizinate injection (MII), DGI, compound glycyrrhizinate tablets (CGT), and diammonium glycyrrhizinate enteric-coated capsules (DGEC) could demonstrate significantly pro-angiogenic effects. Among them, DGI exhibited the strongest activity. Transcriptomic analysis showed that the pro-angiogenic activity of DGI was closely associated with the mTOR signaling pathway. RT-qPCR results indicated that the expression levels of key genes of the mTOR/HIF-α pathway were significantly upregulated in DGI-treated zebrafish relative to the PTK787 group;

**Conclusions:** GA preparations have an obvious pro-angiogenic effect, in which DGI can exert particularly strong activity by activating the mTOR/HIF-1 signaling pathway. This study broadens the clinical application of GA agents and provides new insights into the pharmacodynamics and potential mechanisms of DGI as a candidate agent for treating ischemic diseases

## 1. Introduction

Angiogenesis, the growth of new capillaries from existing capillaries or postcapillary microvessels, plays a pivotal role in a number of biological processes, including embryonic development, wound healing, cardiovascular maturation, and tissue repair [1]. Promoting the angiogenic process is an important strategy in modern medicine for the treatment of ischaemic diseases. For instance, in the case of ischemic heart disease (IHD), also known as coronary heart disease (CHD), the induction of angiogenesis and revascularization is of paramount importance for the effective treatment [2]. The formation of new blood vessels can enhance blood flow in tissues throughout the body, facilitating the delivery of oxygen and nutrients to the infarcted area and its surrounding tissues in a timely manner [3]. Additionally, angiogenesis can inhibit inflammatory and oxidative responses, thereby improving myocardial function. A number of signaling pathways, including VEGF, Notch [4], Wnt/β-catenin [5], Ang1(2)/tie2 [6], PI3K-AKT, and others, are involved in the process of angiogenesis and have an impact on the various stages of neovascularization. Vascular endothelial growth factor (VEGF) is a crucial mediator of numerous signaling pathways, exerting a pivotal influence on the regulation of physiological and pathological angiogenesis [7]. The activation of VEGF has been demonstrated to play an indispensable role to increase vascular permeability, promote endothelial cell proliferation, migration and survival, and regulate the entire process of neovascularization. Research has shown that the phosphatidylinositol 3-kinase (PI3K)/AKT/mammalian target of rapamycin (mTOR) pathway can increase VEGF expression by up-regulating hypoxia-inducible factor 1 alpha (HIF-1α) [8].

Glycyrrhizae radix et rhizoma, as a frequently utilized bulk herb, is the dried roots and rhizomes of Glycyrrhiza uralensis fisch, Glycyrhiza inflata Bat, or Glvcyrrhiza glabra L, Fabaceae [9]. Licorice has a long history of medicinal use, with a wide range of traditional applications to treat respiratory disorders, epilepsy, fever, gastric ulcers, rheumatism, and skin disorders [10]. The principal component of this herb is licorice saponin, also known as GA or glycyrrhizin [11]. GA has a pair of differential isomers, 18α-GA and 18β-GA, due to the difference in the configuration of the chiral carbon atom C-18 of the triterpene saponin. Their hydrolyzed products are 18α-glycyrrhetinic acid and 18β-glycyrrhetinic acid, reapectively [12]. Different configurations of GA also differ in pharmacology, pharmacodynamics, pharmacokinetics and adverse effects [13], Studies have shown that GA has many pharmacological effects, such as protecting hepatocyte membranes, anti-inflammation, antiviral activity, immunomodulation, reducing hepatocyte degeneration and necrosis, reduces gammaglutaminase, inhibiting hepatic collagen fibers proliferation [14], preventing the formation of hepatic fibrosis, promoting bilirubin metabolism, etc.

GA is usually present in the form of a salt when used as a medicine to increase its water solubility [15]. There are many compound preparations of GA in pharmaceutical market, which have been iterated for four times from the original mixed extract of liquorice. The first generation was glycyrrhizin (now unused), the second generation was β-monoammonium glycyrrhizinate salts(e.g., Compound Glycyrrhizin, Compound Monoammonium Glycyrrhizinate Injection, Compound Ammonium Glycyrrhetate and Sodium Chloride Injection, etc.), the third generation was diammonium glycyrrhizinate salts mixed with α- and β-configurations (e.g., DGEC, DGI, etc.), and the fourth generation was magnesium iso-glycyrrhizinate in α-configuration (e.g., MII) [16]. GA preparations are one of the first-line drugs used in anti-inflammatory and hepatoprotective therapy, with the products of diammonium glycyrrhizinate (DG) family as the most commonly used in clinical practice. DG possesses antioxidant activity and has the superior stability and biological activity for the treatment of liver injury than conventional GA. Xie [17] et al. found that DG was able to ameliorate lipopolysaccharide-induced acute lung injury by modulating vascular endothelial barrier function [18]. DGI is an artificial preparation extracted and manufactured from the traditional Chinese medicine liquorice, which plays a key role in anti-inflammatory and hepatoprotective therapy, and is the main pharmacological agent for the treatment of chronic hepatitis B [19]. At present, GA has been found to make the wound dressing together with other chemical substances, such as cumquat glycosides and carvacrol, which exhibited the pro-angiogenic effect to promote wound healing. But the pro-angiogenic activities of GA preparations have not been compared and evaluated specifically, and their mechanism of action is also not clear [20, 21].

The current models for the evaluation of pro-angiogenic activity include the zebrafish model, the rat model, and the Human Umbilical Vein Endothelial Cells (HUVEC) model [22]. Among them, the zebrafish model has excellent biological characteristics, such as small body size, vascular fluorescence, short reproduction cycle, large spawning capacity, low experimental cost, etc., which makes it a very desirable experimental model [23]. In 2002, Lawson and Weinstein constructed the transgenic zebrafish line *Tg (fli-1:EGFP)* with vascular endothelial cells labeled by green fluorescent protein, which has realized the dynamic observation of in vivo vascular development and generating [24]. Transgenic fluorescent labeled zebrafish as the ideal in vivo model for studying metabolic angiogenesis without dissection facilitated activity evaluation related to angiogenesis [25]. And the vascular anatomy of the developing zebrafish embryo has remarkable similarities to that of other vertebrates [26]. PTK787 is a vascular endothelial cell growth factor receptor inhibitor that can inhibit the intersegmental angiogenesis of zebrafish. At present, zebrafish vascular injury model constructed using PTK787 are widely used in activity evaluation and action mechanism research [27, 28].

In this study, we evaluated the pro-angiogenic activity of several common GA preparations on vascular-injured zebrafish to screen the strongly active preparation, and further investigated its pro-angiogenic mechanism of action. Firstly, five GA preparations were selected for pro-angiogenic activity screening, including MII, DGI, CGT, DGEC, and compound monoammonium glycyrrhizinate S for injection (CMGSI). Subsequently, the activity of the represent active preparation DGI was further evaluated and verified. In the end, the potential action mechanism of DGI was investigated for the first time by transcriptomics and RT-qPCR analysis. This work provided experimental bases for the research and development of DGI as a new drug for the prevention and treatment of ischemic diseases. **Figure 1**.

**Figure 1.**
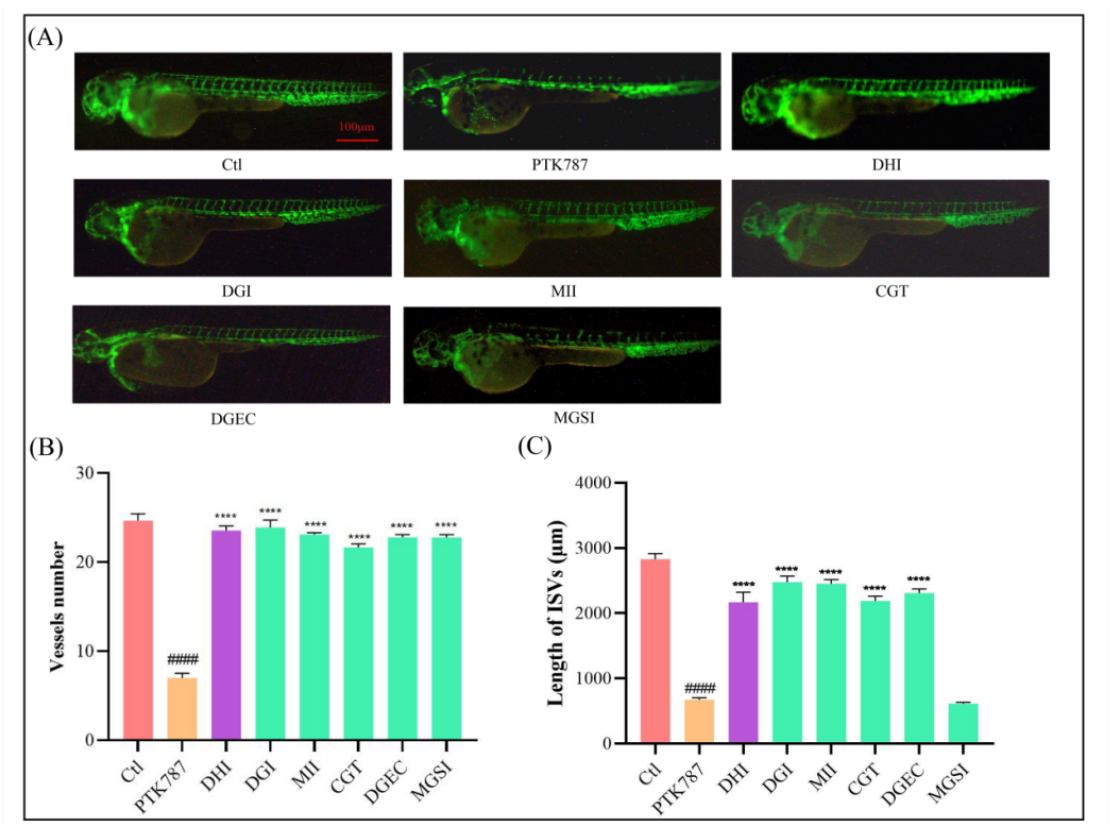
Effects of GA formulations on zebrafish ISV angiogenesis. (**A**) Typical images of zebrafish ISVs (scale: 100 μm). (**B**) Histogram of data analysis of the number of ISVs. (**C**) Histogram of data analysis for length of ISVs. *####p* < 0.0001 compared to the Ctl group, *\*\*\*\*p* < 0.0001, compared to the PTK787 group.

## 2. Materials and Methods

### 2.1. Chemicals and Regents

PTK787 (M18801) purchased from AbMole Biotechnology (Shanghai) Co. Danhong Injection (20230817) purchased from Shandong Danhong Pharmaceutical Co., Diamine glycyrrhizinate injection (21030525) obtained from China Resources Shuanghe Limin Pharmaceutical Co., Magnesium isoglycyrrhizinate (220517104) purchased from Chengtai Tianqing Pharmaceutical Group Co., Compound Glycyrrhizin (21309) obtained from Eisai Pharmaceutical Co., Diamine glycyrrhizinate enteric capsule (220625110) purchased from Chengtai Tianqing Pharmaceutical Group Co., Monoammonium glycyrrhizinate Compound for Injection S (01211002) purchased from Shanxi Puder Pharmaceutical Co., Purified water purchased from Hangzhou Wahaha Group Co., Streptavidin E (516E011) bought from Beijing Soleberg Technology Co., Dimethyl sulfoxide (A100231-0500), Methylene blue (20110520) purchased from Pharmaceutical Group Chemical Reagent Co., Phenylthiourea (PTU) (BCBW4842) purchased from Sigma Corporation USA, Deionized water (FOO20) purchased from Beijing Soleberg Technology Co. The production lot number of FastPure Cell/Tissue Total RNA Isolation Kit V2 is RC112, the production lot number of Hiscripter ?Q RT Super Mix for qPCR (+gDNA wiper) is R223-01, and the production lot number of ChamQ Universal SYBR qPCR Master Mix is Q77; they were purchased from Nanjing Novozymes Bio-technology Co., and RNase-free ddH2O (P071) was bought from Beijing Dingguo Changsheng Biotechnology Co.

### 2.2. Maintenance of zebrafish

The zebrafish strain used in this experiment was *Tg (fli-1:EGFP)* zebrafish, which was provided by the drug screening laboratory in the Institute of Biology, Shandong Academy of Sciences. Zebrafish were cultured according to standard procedures, with male and female zebrafish maintained separately in a recirculating culture system at 28 ± 0.5 °C with a light/dark cycle of 14/10 hours, and fed twice daily. Sexually mature and normally developed zebrafish were selected and placed in the spawning tank at a ratio of 2:2 for male to female. The middle partition of the spawning tank was drawn off at 8:30 on the following day. Zebrafish were stimulated by light to mate and spawn. Two hours later, embryos were collected, washed, and placed in a zebrafish culture solution containing 5 mM NaCl, 0.17 mM KCI, 0.33 mM CaCl_2_, and 0.33 mM MgSO_4_ with the addition of 0.5 mg/L methylene blue. Fish eggs were cultured in a constant temperature light incubator at 28 ± 0.5 °C, and the culture medium was changed every 24 hours. All animal care and experimental procedures were approved by the Experimental Animal Welfare Ethics Committee of the Biology Institute of Shandong Academy of Science.

### 2.3. Proangiogenic activity screening of GA preparations

Healthy transgenic zebrafish *Tg (fli-1:EGFP)* at 24 hpf were selected under Olympus microscope and transferred into 24 well plates. Zebrafish were randomly divided into blank control group (Ctl), model group (0.2 μg·mL^-1^ PTK787), positive control group (0.2 μg·mL^-1^ PTK787 + 9 μL·mL^-1^ DHI), and sample-treated group (0.2 μg·mL^-1^ PTK787 +100 μL·mL^-1^ GA preparations). After administration, Ctl group, PTK787 group, DHI group were placed in a constant temperature incubator at 28 °C for 24 hours. The ISVs of zebrafish in each group were observed under a Zeiss fluorescence microscope. The total length and number of zebrafish ISVs were measured and counted separately using Image-Pro Plus 5.0.

### 2.4 Determination of safe concentrations of DGI

The vascular fluorescent transgenic zebrafish with normal development at 24 hours post fertilization (24 hpf) were put into 24-well plates, and randomly divided into the treatment group of the samples to be tested and the control group, with 10 embryos in each well. Three replicate wells were set up. Zebrafish in the sample groups were treated with different concentrations of DGI at 50, 100, 200, 400, and 800 μg·mL^-1^, respectively. The specific dosage was calculated according to the DG content in DGI preparation. All the above groups of zebrafish were placed in a constant temperature light incubator at 28 ± 5 °C for a total of 24 hours. After cleaning, the survival zebrafish were observed under an Olympus microscope, and the number of zebrafish deaths was counted using the absence of heartbeat as the criterion.

### 2.5 Proangiogenic activity evaluation of DGI on zebrafish ISVs

Except for the test concentrations of DGI, the experimental methods were the same as described in Section 2.3. In this assay, we evaluated the proangiogenic activity of DGI at a gradient concentrations (25, 50, 100 μg·mL^-1^).

### 2.6 Effect of DGI on zebrafish SIVs

Healthy transgenic zebrafish at 72 hpf were randomly transferred into 24-well plates, which were divided into a blank control group, a positive control group (9μg·mL^-1^ DHI), and DGI-treated groups (25, 50, 100 μg·mL^-1^). After 24 hours of drug treatment, SIVs growth in zebrafish were observed and images were captured under a Zeiss fluorescence microscope. SIVs length was measured using Image-Pro Plus 5.0.

### 2.7 mRNA-sequencing and bioinformatics analysis

Samples of Ctl, PTK787, and DGI administration (100 μg·mL^-1^) groups were prepared using the same strategy as it described in “Section 2.3” with 30 embryos in each group. Three replicates of each treatment were subjected to RNA sequencing. Zebrafish were rinsed for three times with purified water separately. RNA was extracted using the TRIzol kit (Invitrogen, US) and stored at 80 °C. The library was subjected to RNA-seq using the llumina Novaseq 6000 sequencing platform to generate 150 bp paired-end sequences. The raw reads in fastq format were processed by fastp software, and cleanreads were obtained after removing low-quality reads for subsequent data analysis. The cleanreads were mapped to the reference genome using software HISAT2. Fragments Per Kilobase Million (FPKM) of each gene was calculated and the read counts of each gene were obtained by HTSeq-count. Principal Component Analysis (PCA) analysis of genes was performed using R (v 3.2.0) as well as mapped to assess sample biological replicates. Differentially expressed genes (DEGs) were analyzed using DESeq2 software with the thresholds of p < 0.05 and foldchange > 2 or foldchange < 0.5. Hierarchical clustering analysis of DEGs was performed using R (v 3.2.0) to demonstrate the expression patterns of genes across groups and samples. Subsequently, enrichment analysis of DEGs based on hypergeometric distribution algorithms such as GO and KEGG Pathway was used to screen for significantly enriched functional entries and pathways. R (v 3.2.0) was used to plot bar charts, enrichment analysis circle plots, etc. Gene set enrichment analysis was performed using GSEA software.

### 2.8 RT-qPCR analysis

According to the experimental method in “Section 2.7”, zebrafish larvae from each group were collected separately for RT-qPCR assay. Total RNA was extracted using RNA extraction kit (Vazyme Biotech Co., Ltd, Nanjing, China), and reverse transcribed into cDNA for qPCR using HiScript II Q RT SuperMix. Subsequently, the cDNA was used as a template for the reaction using ChamQ Universal SYBR qPCR Master Mix (Vazyme Biotech Co., Ltd, Nanjing, China). The reaction conditions were 1 cycle of pre-denaturation at 90 °C for 30 s, followed by 40 cycles of denaturation at 95 °C for 15 s and annealing at 60 °C for 30 s, and 1 cycle at 60 °C for 60 s and 95 °C for 15 s. The experimental data were analyzed by the 2^-ΔΔCt^ method using β-actin as an internal reference to calculate the expression levels of each gene in zebrafish from each group. BioSune Biotechnology (Shanghai) Co, Ltd. synthesized the primers of all genes and their sequences are shown in table 1.

**Table 1.**
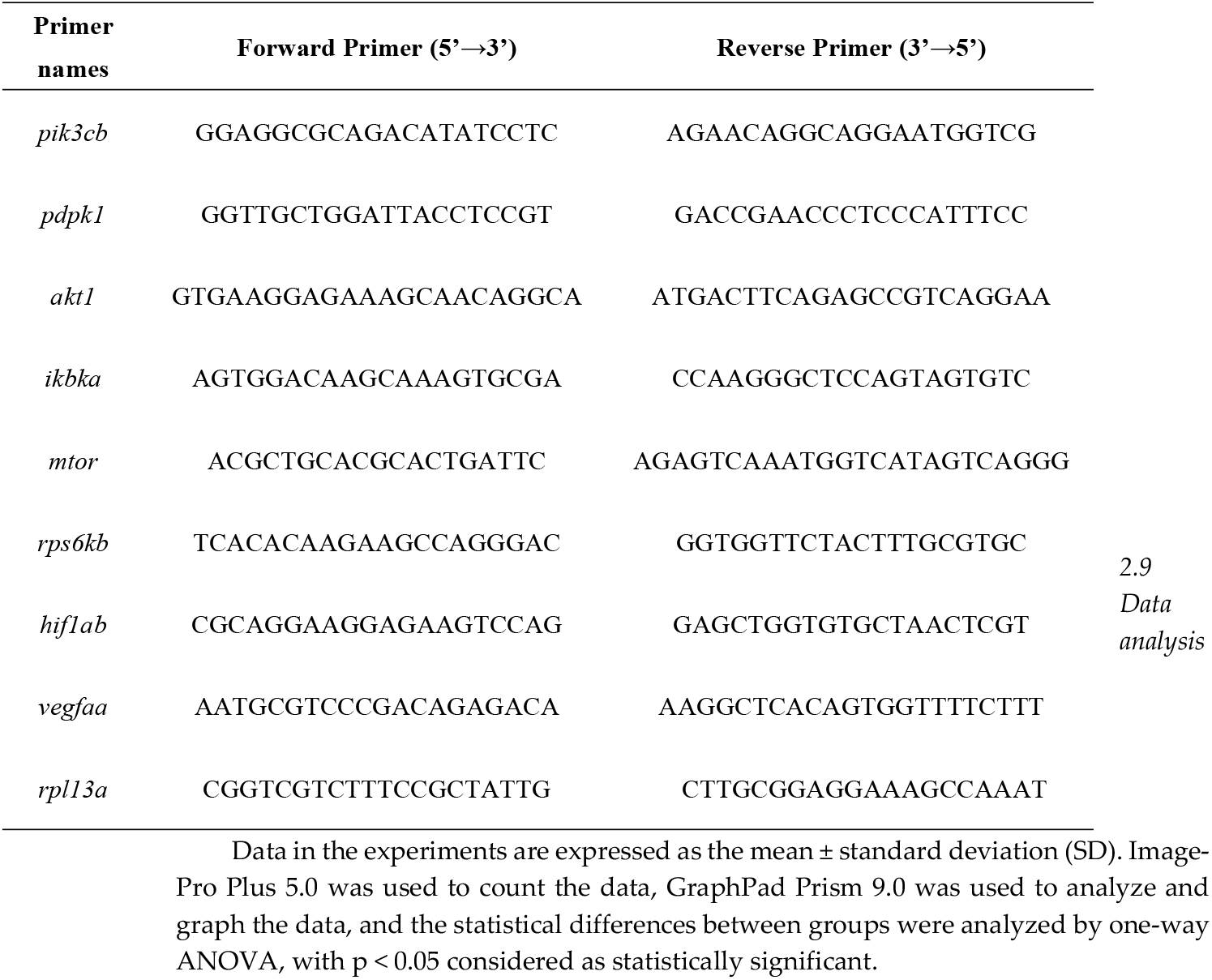
Primer sequences

Data in the experiments are expressed as the mean ± standard deviation (SD). Image-Pro Plus 5.0 was used to count the data, GraphPad Prism 9.0 was used to analyze and graph the data, and the statistical differences between groups were analyzed by one-way ANOVA, with p < 0.05 considered as statistically significant.

## 3. Results

### 3.1. Screening of GA compound preparations with pro-angiogenic activity

Zebrafish *Tg (fli-1:EGFP)* was utilized to study the effects of different GA compound preparations on the angiogenesis in zebrafish (Figure 1). Compared with the blank control group, both of the length and the number of zebrafish ISVs in the PTK787 model group were significantly reduced indicating that we successfully constructed the vascular injury model in zebrafish using PTK787. Compared with the PTK787-induced model group, the length and the number of zebrafish ISVs were significantly increased in the sample groups treated with DGI, MII, CGT, and DGEC, while those indexes didn’t change obviously in CMGSI treatment group at the experimental concentration. This result indicated the marked pro-angiogenic activity of DGI, MII, CGT, and DGEC. Among them, DGI was the most active preparations.

### 3.2. Safety concentration measurement for DGI

Before the activity evaluation, the safety dose of DGI to zebrafish was investigated first. The result showed that zebrafish larvae were normally developed in treated groups with all of concentration groups and no individual died in this experiment (Figure 2). Hence, DGI was nontoxic to zebrafish at the concentration of less than 800 μg·mL^-1^. According to the conversion method of dosage of administration from zebrafish to human (Patent Application Publication No.: CN113496071A; Application No. 202010256107.8), the concentration of 800 μg·mL^-1^ is equivalent to 4 g of drug per day for humans, which suggested the safety of DGI to zebrafish. In the subsequent experiments, 25, 50, and 100 μg·mL^-1^ were selected as the experimental concentrations to evaluate the activity of DGI.

**Figure 2.**
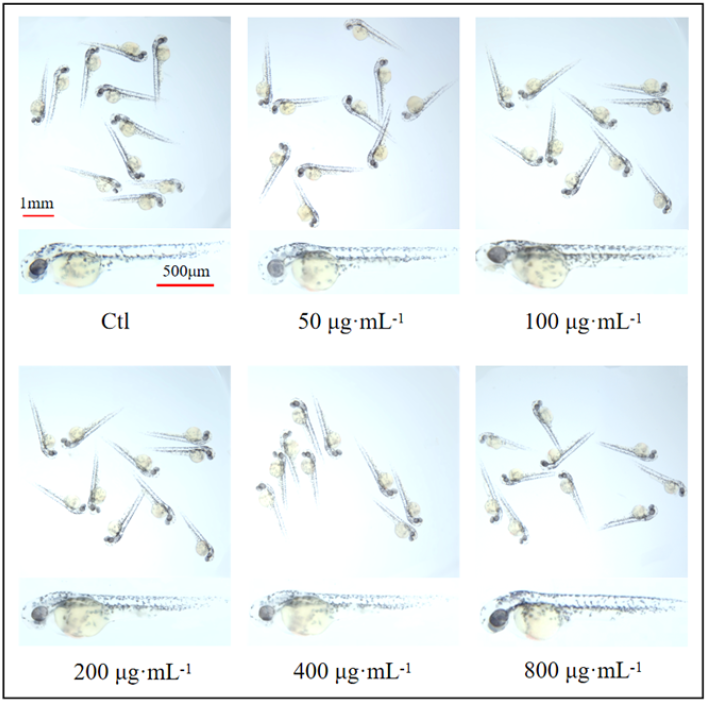
Examination of the safe dose of DGI to zebrafish larvae (scale: 1 mm, 500 μm).

### 3.3 Pro-angiogenic effect of DGI on zebrafish ISVs

The result of the effect evaluation of different concentrations of DGI (25, 50, 100 μ g·mL-1) on the ISVs in zebrafish (Figure 3), indicated that DGI can significantly reverse the decrease in the length and number of ISVs in zebrafish caused by PTK787 in a concentration-dependent manner.

**Figure 3.**
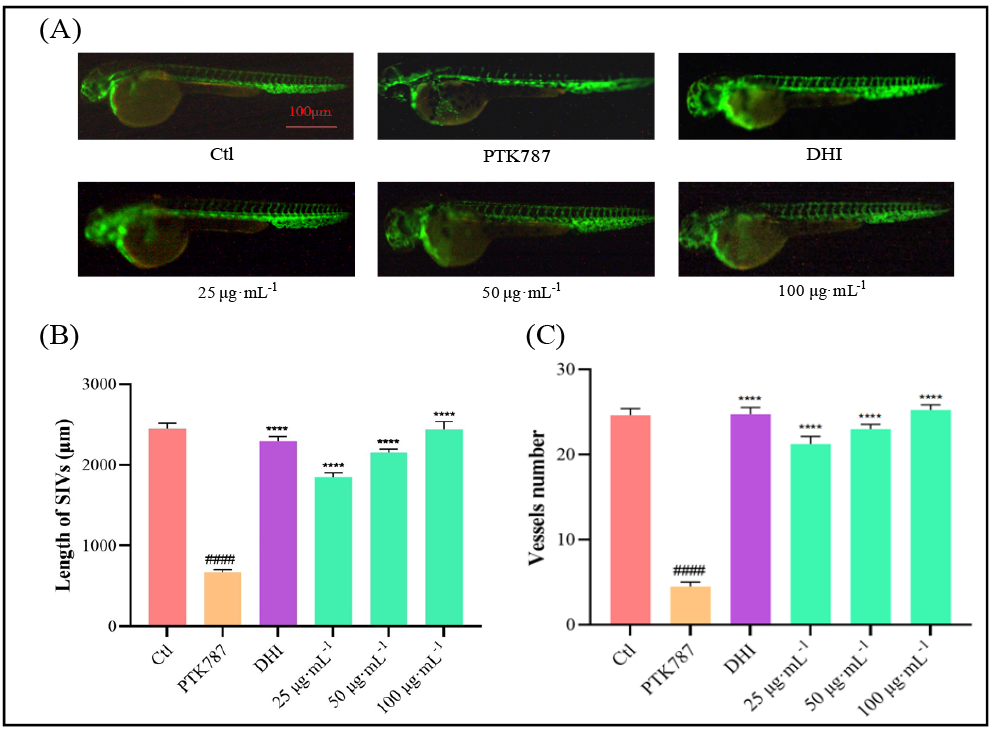
Pro-angiogenic effects of DGI on SIVs in zebrafish. (A) Typical images of zebrafish ISVs (scale: 100 μm). (B) Histogram of data analysis of the length of ISVs. (C) Histogram of data analysis for number of ISVs. *####p* < 0.0001 compared to the Ctl group, *\*\*\*\*p* < 0.0001, compared to the PTK787 group.

### 3.4. Pro-angiogenic effect of DGI on zebrafish SIVs

As shown in Figure 4, compared with the Ctl group, DGI at all the treatment concentrations could significantly increase the length of zebrafish SIVs, indicating a good activity for DHI to promote the growth of zebrafish SIVs.

**Figure 4.**
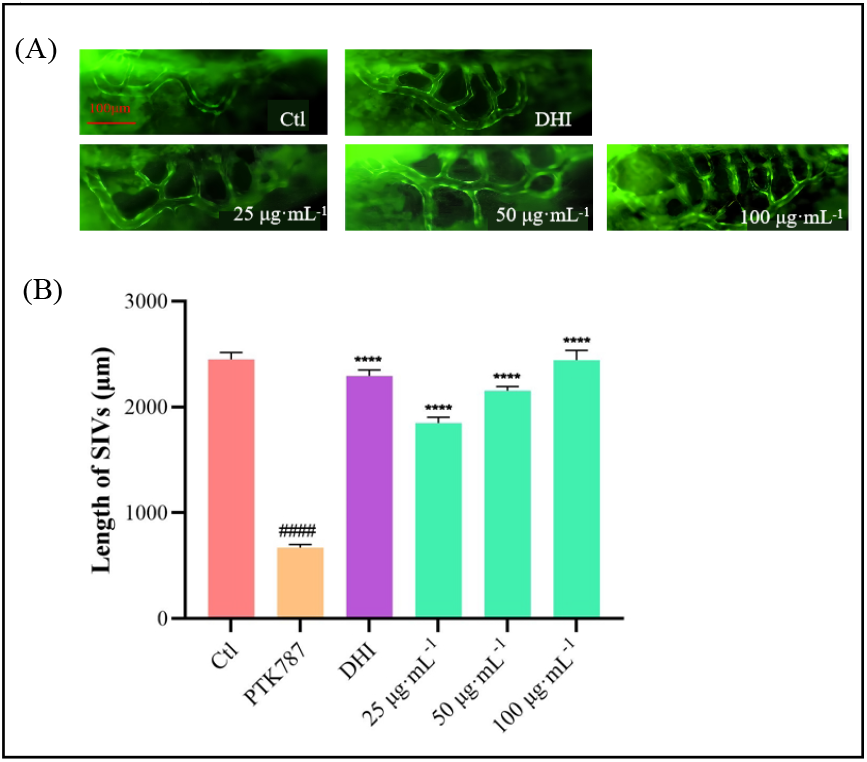
Pro-angiogenic effects of DGI on SIVs in zebrafish. (A) Typical images of zebrafish ISV (scale: 100 μm). (B) Histogram of data analysis of the length of ISVs. (C) Histogram of data analysis for number of ISVs. ####p < 0.0001 compared to the Ctl group, ^****^p < 0.0001, compared to the PTK787 group.

### 3.5. mRNA-sequencing and bioinformatics analysis

#### 3.5.1 Differential gene analysis

The mRNA profiles of the Ctl group, the PTK787 group and the DGI administration group were determined through transcriptome sequencing. Principal component analysis (PCA) was used to assess the biological reproducibility of the samples within the groups and the differences between the samples. Figure 5 (A) illustrates a notable disparity between the three groups, substantiating the efficacy of PTK787 modelling and DGI activity and the reliability of the analytical approach. The volcano plot Figure 5 (B) visualizes the distribution of DEGs for the two compared combinations. Compared with the Ctl group, the PTK787 group had 243 significant DEGs, including 155 up-regulated and 88 down-regulated genes. Compared with the PTK787 group, the DGI group had 54 significant DEGs, including 22 up-regulated and 33 down-regulated genes. A total of 24 co-DEGs in total were obtained between the comparisons of PTK787 group vs Ctl group and DHI group vs PTK787 group Figure 5 (C). 15 of these DEGs were upregulated in the comparison of PTK787 group vs Ctl group and downregulated in the comparison of DHI group vs PTK787 group. Other 9 DEGs showed the opposite change.

**Figure 5.**
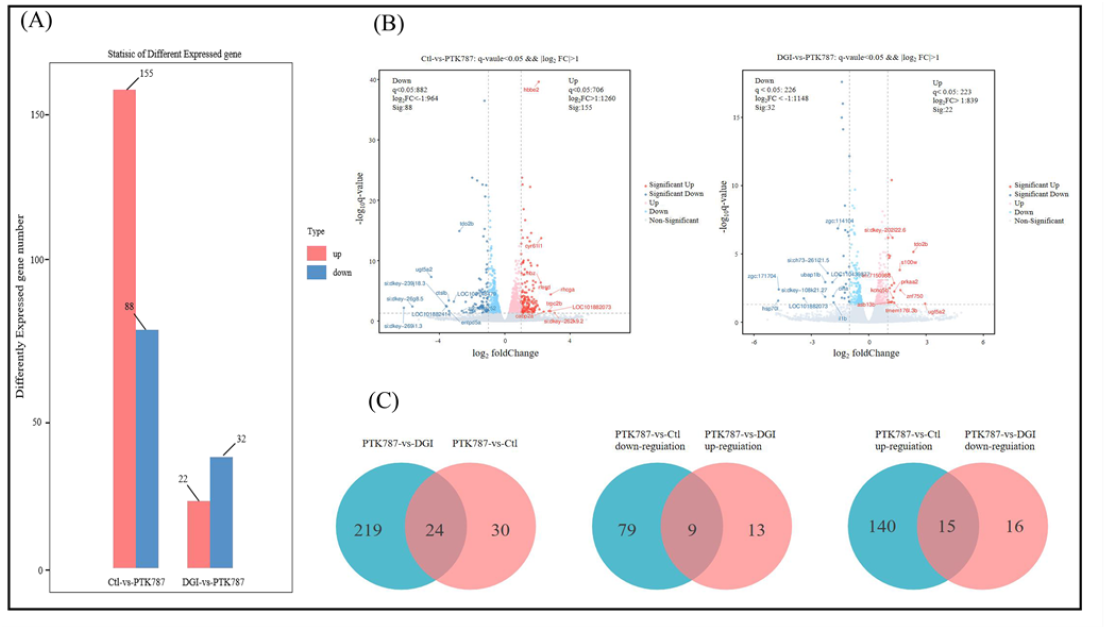
Analysis of DEGs before and after DGI administration in zebrafish. (A) Histogram of DGEs statistics. (B) DGEs volcano map. (C) Venn diagram of DGEs.

#### 3.5.2 GO enrichment analysis

Then, the DEGs were employed for next analysis of GO functional enrichment. GO is a comprehensive database describing gene functions, which includes three parts: Biological Process (BP), Cellular Component (CC), and Molecular Function (CF). As can be seen in Figure 6 (A). altered BP-related items in the DGI-treated group compared with the PTK787 group included nitric oxide biosynthetic process, tryptophan catabolic process, etc. For the CC-related parts, DGI mainly acted on SOSS complex, vesicle membrane, intermediate filament, etc. As for the MF-related items, DGI mainly regulated protein folding chaperone, collagen binding, promoter-specific chromatin binding, and so on. As can be seen in Figure 6 (B). altered BP-related items in the Ctl group compared with the PTK787 group included erythrocyte development, GPI anchor biosynthetic process, ventricular cardiac muscle cell develop etc. For the CC-related parts, DGI mainly acted on oxygen transport basolateral plasma membrane, integrator complex, endosome membrane, etc. As for the MF-related items, DGI mainly regulated metal ion binding, L-ascorbic acid binding, cysteine-type endopeptidase activator, and so on.

**Figure 6.**
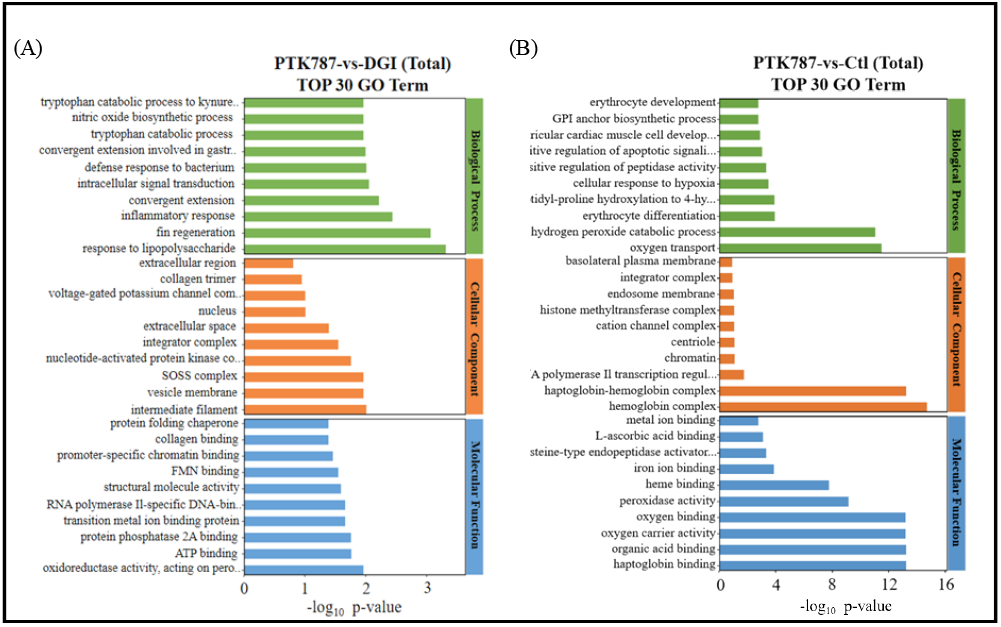
Results of GO functional enrichment analysis (Top 30 terms) for DEGs between PTK787 and DGI groups (A), and between Ctl and PTK787 groups(B).

#### 3.5.3 KEGG enrichment analysis

We performed KEGG enrichment analysis on the screened DEGs. From the comparison between the model group and control group, 68 pathways were enriched, while for the comparison between the DGI-treated group and model group, 43 pathways were enriched. Top 20 pathways by KEGG analysis were shown in Figure 7, including mTOR signaling pathway, MAPK signaling pathway, Wnt signaling pathway, and some metabolism-related pathways, and so on. Among them, the mTOR signaling pathway is particularly outstanding in both of the two comparison pairs of the model group vs control group and the DGI-treated group vs model group, and closely related to angiogenesis. So in this study, we selected key genes of the mTOR signaling pathway for next RT-qPCR validation.

**Figure 7.**
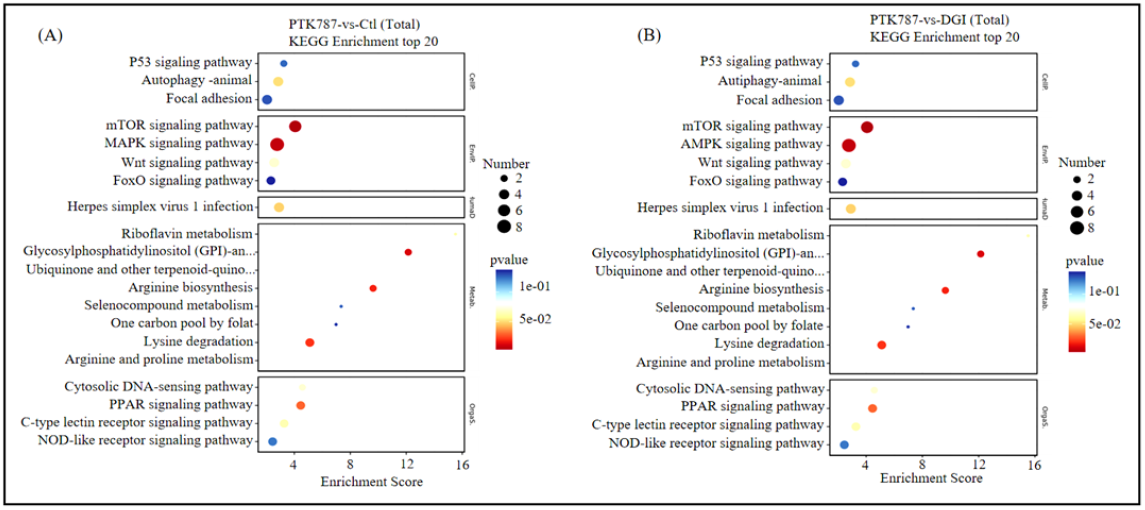
Results of KEGG pathway enrichment analysis (Top 20 pathways) for DEGs between PTK787 and Ctl groups (A), and between DGI treatment and PTK787 groups (B).

### 3.6 RT-qPCR analysis

We detected the expression levels of the mTOR signaling pathway and its downstream pathway/HIF-1 pathway genes *pik3cb, pdpk1, akt1, ikbka, mtor, rps6kb, hif1ab, vegfaa* in zebrafish before and after DGI administration. The results showed that the PTK787 group significantly down-regulated the expression of the above genes compared to those in the Ctl group. Compared with the PTK787 group, the expression levels of the above genes were significantly enhanced by DGI treatment Figure 8. The above results suggested that DGI could activate mTOR signaling pathways to promote the angiogenesis in zebrafish.

**Figure 8.**
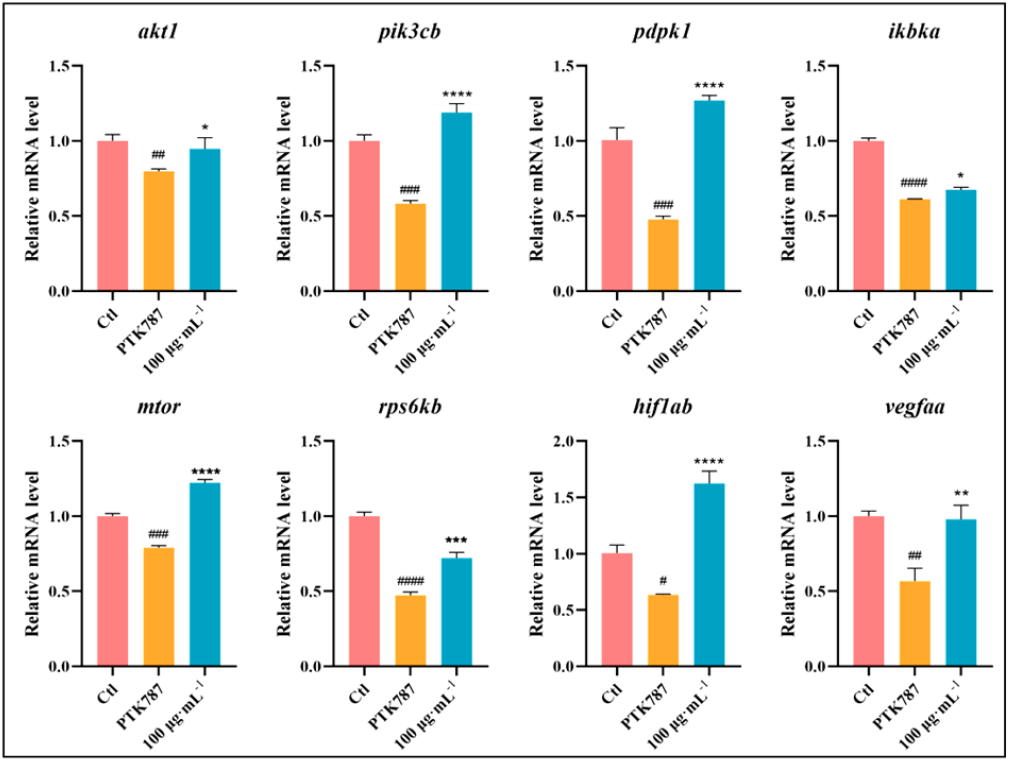
Effects of DGI on the expressions of key genes in zebrafish. *####p* < 0.0001, *###p* < 0.001, *##p* < 0.01, #p < 0.05 compared to the Ctl group; *\*\*\*\*p* < 0.0001, *\*\*\*p* < 0.001, *\*\*p* < 0.01, and *\*p* < 0.05 compared to the PTK787 group.

**Figure 9.**
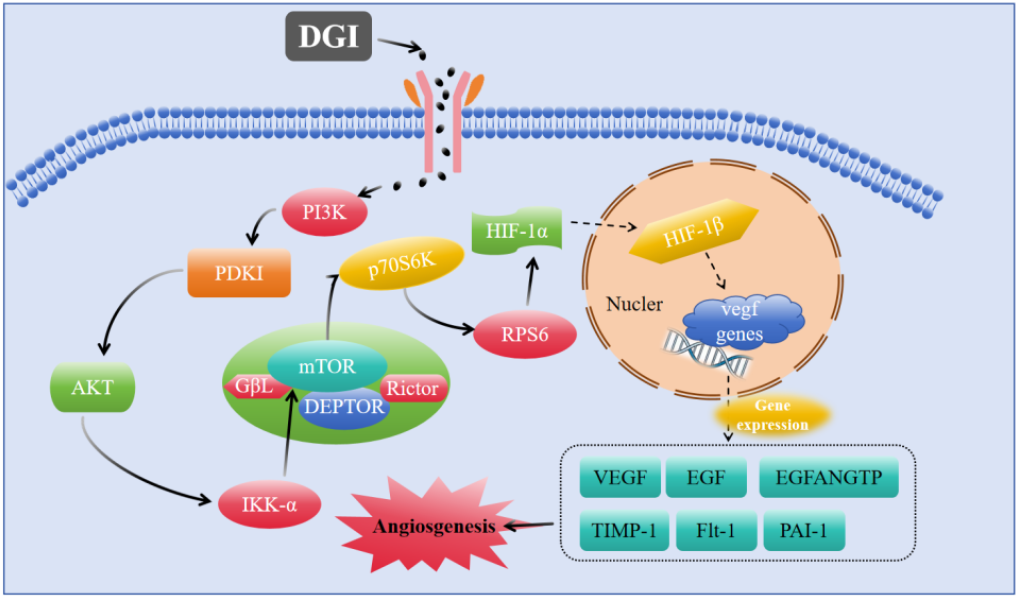
Schematic diagram of the potential mechanism of DGI pro-angiogenesis.

## 4. Discussion

GA preparations are commonly used in clinical anti-inflammatory and liver protection drugs [29]. However, the pro-angiogenic activity and mechanism of GA preparations have not been studied specifically. In this study, we initially screened a number of effective GA preparations with the pro-angiogenic activity using a vascular injury zebrafish model induced by PTK787. Therein, preparation DGI exhibited the strongest activity, which can significantly improve the angiogenesis of zebrafish ISVs and SIVs at a concentration-dependent manner. Furthermore, the mechanism of action for DGI was investigated using transcriptomics analysis and experimental validation by RT-qPCR. Our results indicated that DGI promoted angiogenesis in zebrafish mainly via regulating mTOR signaling pathways.

To further explore the potential mechanisms by which DGI promotes angiogenesis, we used transcriptomics to speculate on the signaling pathways affected by DGI. By transcriptomics analysis, 243 and 54 significant DEGs were obtained from the comparisons of PTK787 group vs Ctl group and DGI group vs PTK787 group, respectively. On based of these DEGs, we performed GO and KEGG enrichment analysis and found that mTOR signaling pathway was a key pathway for DGI to exert pro-angiogenic effect. mTOR signaling pathway is a classical signaling pathway to regulate the cell cycle, autophagy, apoptosis, proliferation, and other physiological processes [30]. Alterations in the genes associated with this pathway are closely linked to the treatment of cardiovascular diseases [31], tumorigenesis [32], immune disease [34], and so on. As a crucial substrate of Akt, mTOR is a serine/threonine kinase, which regulates various cellular life activities by modulating protein synthesis, glucose metabolism, and lipid metabolism, thereby playing an essential role in promoting cell proliferation and growth [34]. Akt is able to activate mTOR in two ways, including by phosphorylating mTOR directly or by inactivating Tuberous Sclerosis Complex 2 (TSC2), which maintains the GTP-bound state of the Ras homolog enriched in the brain (Rheb), furtherly enhancing mTOR activation [35]. Eukaryotic translation initiation factor 4E-binding protein1 (4E-BP1) and p70 ribosomal protein S6 kinase (p70S6K) are two important effector molecules downstream of mTOR. mTOR activation can phosphorylate both 4E-BPl and p70S6K.

Hypoxia inducible factor (HIF), located downstream of p70S6K, is a heterodimer composed of HIF-1α and HIF-1β [36]. Studies found that the hypoxic microenvironment after traumatic surface formation is a key initiator of angiogenesis [37]. Activation of the HIF-1 signaling pathway is the main mechanism for tissue and cellular responses to hypoxia, and the HIF-1 signaling pathway is closely related to angiogenesis. HIF-1α forms a heterodimer with HIF-1β through post-translational modification and binds to hypoxia response elements (HREs) on the promoters of various genes, promoting the transcription of the target gene vegf, which in turn leads to the rapid proliferation of endothelial cells to promote angiogenesis [38]. The mTOR signaling pathway and its downstream HIF-1 signaling pathway play important roles in pro-angiogenesis. In the mTOR/HIF-1 signaling pathway, the activation of the genes *pik3cb, pdpk1, akt1, ikbka, mtor, rps6kb, hif1ab*, and *vegfaa* showed a cascade reaction, starting from *pik3cb* and then activating the other genes, one by one in turn. When mtor is activated by a series of upstream genes, its gene expression product mTOR phosphorylates the expression product of genes rps6kb and p70S6K. It then activates the expression of the downstream gene hif1ab and the interaction between HIF-1α and HIF-1β proteins, which ultimately promotes the expression of the target gene vegfaa. The potential mechanisms are shown in Figure 10. Expression product VEGF of gene vegfaa can promote the growth and migration of vascular endothelial cells and accelerate angiogenesis. In this study, we validated the effects of DGI treatment on the above related gene expressions in zebrafish by RT-qPCR. Results showed that the expression levels of genes pik3cb, pdpk1, akt1, ikbka, mtor, rps6kb, hif1ab, and vegfaa were significantly up-regulated in the zebrafish after DGI administration relative to the PTK787 group, demonstrating that DGI was able to activate the mTOR/HIF-1 signaling pathway to exert pro-angiogenic effects.

In this study, we first discovered the in vivo pro-angiogenic activities of GA-based agents DGI, MII, CGT, and DGEC using zebrafish model, and revealed the underlying mechanism of DGI by transcriptomics and RT-qPCR analysis. Our research suggests that DGI. However, there are still some limitations: 1) the specific targets of DGI need to be verified thoroughly by Western blotting and gene-editing experiments, 2) other GA formulas with pro-angiogenic activity need to be further investigated for their mechanisms.

## 5. Conclusions

In this study, we compared the pro-angiogenic activities of five commonly used GA-based agents for the first time, namely DGI, MII, CGT, DGEC, and CMGSI, using zebrafish vascular injury model, and screened out DGI with the most pronounced activity, then further elucidated the mechanism of the pro-angiogenic effect of DGI by transcriptomics and RT-qPCR analyses. The results indicated that DGI can promote angiogenesis in zebrafish by activating the mTOR/HIF-1 signaling pathway. This paper provides an experimental basis for the study and development of DGI as a new drug to treat ischaemic diseases.

## Author Contributions

Conceptualization, S.Y. and X.L.; methodology, S.Y., J.Y. and F.S.; software, W.S. and K.L.; validation, W. L.; formal analysis, X.L. and S.Y.; investigation, S.Y. and H.L.; data curation, S.Y. ; writing—original draft preparation, S.Y. ; writing—review and editing, S.Y. and X.Z.; visualization, W.L.; supervision, X.L. and K.L.; project administration, S.Y. and K.L.; All authors have read and agreed to the published version of the manuscript.

## Data Availability Statement

The data presented in this study are available on request from the corresponding author due to the original contributions presented in the study are included in the article.

